# High density single-unit human cortical recordings using the Neuropixels probe

**DOI:** 10.1101/2021.12.29.474489

**Authors:** JE Chung, KK Sellers, MK Leonard, L Gwilliams, D Xu, M Dougherty, V Kharazia, M Welkenhuysen, B Dutta, EF Chang

**Affiliations:** Department of Neurological Surgery, University of California, San Francisco, California 94143; IMEC, Leuven, Belgium

## Abstract

A fundamental unit of neural computation is the action potential. While significant advances have been made in the ability to sample action potentials of large numbers of individual neurons in animal models, translation of these methodologies to humans has been lacking due to clinical time constraints, electrical noise in the operating room, and reliability of the methodology. Here we present a reliable method for intraoperative recording of dozens of neurons in humans using the Neuropixels probe, yielding up to ∼100 simultaneously-recorded single-units (n=596 across 11 recordings in 8 participants). Most single-units were active within 1 minute of reaching target depth, compatible with clinical time constraints. Cell pairs active close in time were spatially closer in most recordings, demonstrating the power to resolve complex cortical dynamics. Altogether, this approach provides access to population single-unit activity across the depth of human neocortex at scales previously only accessible in animal models.

**Highlights:**

- Single units in 8 patients, yielding 596 putative single units across 11 recordings
- The majority of putative neurons fire at least 1 spike within one minute after reaching target depth
- Putative neurons take longer to fire at least one spike in anesthetized compared to awake participants
- Cell-pairs that fire action potentials close in time are spatially closer together than those fire further apart in time

## Introduction

The study of human neural processing would benefit from our ability to observe the dynamics of the underlying substrate, action potentials of single neurons and the networks they form. This, in turn, requires electrodes to be implanted directly into the brain (Cash and Hochberg, 2015; Hong and Lieber, 2019). Direct access to the intact human brain for single-neuron recordings can either be done acutely in the intraoperative setting, semi-chronically for epilepsy monitoring, or chronically - though for relatively narrow clinical indications and correspondingly diminutive patient populations. Intraoperative single-neuron recordings have potential access to far larger patient populations but are severely limited in the number of simultaneously recorded single neurons. The most often utilized sharp metal microelectrodes typically yield a single neuron at a time (Benazzouz et al., 2002; Guridi et al., 2000; Hutchison et al., 1998). Microwire-based research-focused intraoperative technologies such as the Behnke-Fried electrode can yield up to 2.7 neurons per 8-microwire depth array on average, with 1-2 neurons being the most common yield (Misra et al., 2014). For semi-chronic or chronic studies, the Utah electrode array, a 96-channel ‘bed-of-nails’ array (Maynard et al., 1997; Nordhausen et al., 1994) has been utilized for brain machine interface (Ajiboye et al., 2017; Collinger et al., 2013; Gilja et al., 2015; Hochberg et al., 2012) or memory studies (Vaz et al., 2020), but lacks the ability to sample across the cortical depth and depends upon far smaller patient populations making patient heterogeneity a significant barrier. An ideal technology for studying human neural computation would have access to large numbers of potential human participants and simultaneously record hundreds of neurons.

The past decade has seen the development of an impressive armament of new tools for neuroscientists to understand the structure and function of the brain in animal models, many of which are designed specifically for simultaneous recording from large numbers of single neurons (Chen and Pesaran, 2021). Many of these advances have been driven by high-density microfabricated electrode arrays and associated developments in integration (Berényi et al., 2014; Chung et al., 2019; Jun et al., 2017; Musk and Neuralink, 2019; Shobe et al., 2015; Steinmetz et al., 2021), thereby providing access to spatial and temporal scales required to understanding information processing across the cortical microcircuit (Larkum, 2013). However, translation of methods for recording fundamental computational units (e.g., single neurons, ensembles) has proven extremely difficult. This has led to an increasing gap between what can be learned in animal models and how these detailed principles apply to the human brain.

Here, we present a methodology that utilizes the Neuropixels probe (Dutta et al., 2019; Jun et al., 2017; Steinmetz et al., 2018) and addresses limitations of patient populations and simultaneously recorded neuron yields while recording across the cortical depth. The Neuropixel probe combines advanced complementary metal-oxide-semiconductor (CMOS) technology with integrated amplification, digitization, and multiplexing circuits together on the same device for large-scale, high-density recordings with streamlined data transfer without bulky cabling or connectorization. This probe contains 960 12μm x 12μm contacts with 384 selectable recording channels, arranged along a 10mm length shank in a 4-column checkerboard pattern, with 16μm pitch across columns and 20um pitch across rows. The Neuropixels probe has seen rapid adoption across multiple animal models in the neuroscience community(van Daal et al., 2021; Gaucher et al., 2020; Juavinett et al., 2019; Jun et al., 2017; Trautmann et al., 2019).

Here, and in simultaneously-completed early experience (Paulk et al., 2021), the Neuropixels probe shows potential to access dozens of simultaneously-recorded single-neurons in human participants after addressing translational issues including the safety and sterilization concerns, and electrical noise mitigation in the intraoperative setting. In this work we report on 596 putative single-units with cluster quality metrics from 11 recordings across 8 of our first 15 consecutively tested human participants, all of whom undergoing craniotomy for a variety of clinical indications. Here we demonstrate the method’s potential for access to broad participant populations, dozens of simultaneously-recorded single-neurons, feasibility for intraoperative use, and access to a variety of target locations.

## Methods

### Probe preoperative preparation and sterilization

Neuropixels 1.0 NHP-short probes, with 10mm long shanks and metal caps, were used for all recordings. These probes are adapted and modified versions of the Neuropixels 1.0 rodent probes (Jun et al., 2017) that are electronically identical, with the notable difference of increased shank thickness from 24μm to 97μm (Paulk et al., 2021). This increased thickness allowed for tolerance of greater mechanical forces for larger brain recordings. Needle electrodes were soldered separately to contacts on the probe flex-interconnect to serve as ground and reference. Probes with soldered needle electrodes, headstages, interface cables, and probe mount with metal cap dovetail were all sterilized using ethylene oxide prior to use. Probes were not reused across participants.

### Participant and recording site selection

The experimental protocol was approved by the UCSF Institutional Review Board. Fifteen participants were included in the study (Table 1), and all provided written informed consent. Eleven participants had medically refractory seizures from temporal lobe epilepsy, requiring surgical management during which the anterior temporal lobe was resected from a lateral approach. Two participants had a tumor and two had a cavernous malformation with surrounding epileptogenic tissue that was resected. In some cases, clinical grid and strip ECoG arrays were used as part of routine clinical care for intraoperative monitoring. Participants were either under general anesthesia or under sedation (awake or asleep, Table 1) during the recordings according to clinical need (e.g. intraoperative stimulation mapping procedures). For all cases using general anesthesia, a combination of dexmedetomidine, propofol, and fentanyl or remifentanil were administered. In cases where recording was done in the awake or asleep state, a combination of dexmedetomidine and remifentanil was administered, with propofol sometimes being used during initial induction.

In all cases, the insertion of Neuropixels probes was targeted to cortical tissue that was destined for resection in the same procedure based on clinical criteria. In cases where a tumor or cavernoma was resected, the recording was targeted to radiographically normal tissue (no T2 hyperintensity) which would require resection as part of the transcortical approach. In all cases the recorded tissue appeared normal intraoperatively. The specific sites were selected to be the crown of surface gyri which allowed for direct visualization and monitoring of the insertion and penetration through cortical layers. See below for post-hoc insertion localization.

### Probe positioning and insertion

Each patient was secured with a Mayfield skull clamp with position depending on clinical indication. In turn, a series of clamps was connected to the skull clamp and finally to a micromanipulator which held the Neuropixels probe assembly. All equipment was sterilized according to standard protocols using either Sterrad or ethylene oxide sterilization. The probe was positioned above the target insertion site, and the Neuropixels probe was lowered using the micromanipulator to a target depth of 6 to 8mm from the brain surface. Insertion trajectory was perpendicular to the surface. Insertion locations were estimated through a combination of intraoperative navigation, intraoperative photos taken during the surgery, and histology when available. In some cases, a sharp piotomy was performed at the site of insertion, which reduced the risk for probe fracture. Recordings were started prior to insertion (Supplemental Movie 1).

### Post-hoc insertion localization

Localization of the probe insertion site was done using preoperative MRI with fiducial markers and intraoperative neuronavigation (Brainlab, Munich, Germany), which together have an average target error of 2.49 mm (Mongen and Willems, 2019). Beyond this localization, intraoperative photos were used to supplement intraoperative navigation. In cases where grid and depth electrodes were implanted prior to recording, these images and combined reconstruction were also used to further supplement insertion localization.

### Recording

Data were collected using a custom-constructed rig including a Windows machine, PXI chassis (NI PXIe-1071), PXI Multifunction I/O Module (NI PXIe-6341), NI SHC68-68-EPM shielded cable, Neuropixels PXIe Acquisition Module, and NI BNC-2110 Connector Block with SpikeGLX 3.0 (http://billkarsh.github.io/SpikeGLX/) acquisition software. In some experiments, speakers were used to present auditory stimuli and a microphone was used for recording; these signals were also acquired as analog inputs synchronized with the neural data. The Neuropixels probe was configured to acquire from 384 channels in a ‘long column’ layout, providing the greatest possible depth span of recording while acquiring from a contact at each depth. Total recording span was 7.67mm. Ground and reference needle electrodes were inserted into the scalp adjacent to the craniotomy. AP gain was 500 and LF gain was 250. During data acquisition, all non-essential equipment in the operating room was unplugged or run using battery power in order to reduce electrical noise. In particular, the operative table, electrocautery, and electronics associated with intravenous lines should be on direct current, including syringe pumps and blood/fluid warmers. Other environmental sources of noise included neuromonitoring and the Brainlab neuronavigation system.

### Spike sorting

Kilosort 2.5 (Pachitariu et al., 2016) was used for spike sorting on a server with an NVIDIA GPU, CUDA, and Matlab installed. Sorting was conducted on data after final recording depth was achieved. Following automated spike sorting, Phy (Rossant et al., 2016) was used for manual curation of clusters by an experienced electrophysiologist (KKS).

### Cluster metrics

Amplitude histogram truncation was calculated using code based upon SpikeInterface (Buccino et al., 2020). This metric provides an estimate of the miss rate due to spike amplitude falling below the detection threshold, based upon the amplitude histogram. The empirical amplitude histogram is modeled using a gaussian, with the fraction of the estimated events below the detection amplitude of the model being reported. Higher proportion of amplitude histogram truncation reflects a higher estimated false negative rate and lower cluster quality.

The proportion of events violating the refractory period (Hill et al., 2011) was calculated using code based upon SpikeInterface (Buccino et al., 2020). This metric reports the number of events in a cluster below an inter-spike-interval threshold of 1 ms. Higher proportion of events that violate the refractory period reflects a higher estimated false positive rate and lower cluster quality.

D-prime (Hill et al., 2011) is a linear discriminant analysis-based metric to quantify isolation distance from other clusters. Higher D-prime reflects higher cluster quality and lower estimated false positive rates.

Isolation score (Chung et al., 2017) is a nearest-neighbor based metric which reports the proportion of spikes whose nearest neighbor in feature space comes from the same cluster. Higher isolation reflects higher cluster quality and lower estimated false positive rate.

### Cell-pair coordination index

Cell-pair coordination index is a cell-pair metric to capture the relative coordinated activity (within 50ms) compared to activity seen at slightly longer timescales (50 to 100ms). Single-unit pair firing relationships were first quantified using cross-correlograms. For each cross correlogram that contained at least 100 events within a 1 second window (±0.5s), the sum of events was calculated in three different windows: from -50 to 50ms (s0), -100 to -50ms (s1), and 50 to 100ms (s2).

The cell coordination index (CCI) was then defined as: 

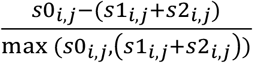

As such, a more positive CCI reflects higher cell-pair activity from -50 to 50 ms compared to - 100 to -50 ms and 50 to 100 ms, while a negative CCI corresponds to higher cell pair activity seen combined in -100 to -50 ms and 50 to 100 ms compared to the -50 to 50 ms window.

### Histology

All tissue that had a device inserted was previously determined to be bound for resection. Despite this, in some cases the tissue could not be safely resected en bloc due to proximity to critical vascular structures, and as such, histology was not performed on all insertion sites. In cases where en bloc resection of the insertion site was possible, the tissue was drop-fixed in ice-cold formalin before being cryoprotected and sectioned into 50μm thick sections. Tissue was then stained with luxol fast blue and neutral red.

## Results

Intraoperative recordings utilizing the Neuropixels probe were attempted in 15 participants (Table 1). A Greenberg C-clamp, articulating bar, 3 ball-joint arm, and micromanipulator were attached to the Mayfield head clamp in series, allowing for gross targeting of the insertion location. The probe was then removed from custom-designed holders used during sterilization and transport and mounted to the micromanipulator (Figure 1A). Targeted regions were in the right or left superior temporal gyrus in 5 of 8 participants with putative single units, right middle frontal gyrus in one case (Figure 1B, Supplemental Figure 1), right ventral motor cortex in one case, and left angular gyrus in one case, dependent solely on the resection plan (Table 1). The insertion locations were at the center of a gyrus, away from surface vasculature, that was going to be resected in the same surgical procedure. In some cases, a piotomy was performed to allow for easier probe penetration. The probe was then inserted to a targeted depth of 7mm (to span the full depth of the cortex and also record channels at the brain/air boundary for movement tracking) (Supplemental video 1). In some cases, the insertion sites were far enough away from local vasculature to allow for safe en bloc resection and subsequent histology (Figure 1C). We found reducing AC noise was key for achieving required signal to noise in the intraoperative environment. Furthermore, reducing positive end expiratory pressure for intubated participants helped reduce the amplitude of brain pulsation reflected in recordings caused by both cardiac and respiratory cycles.

**Figure 1:**
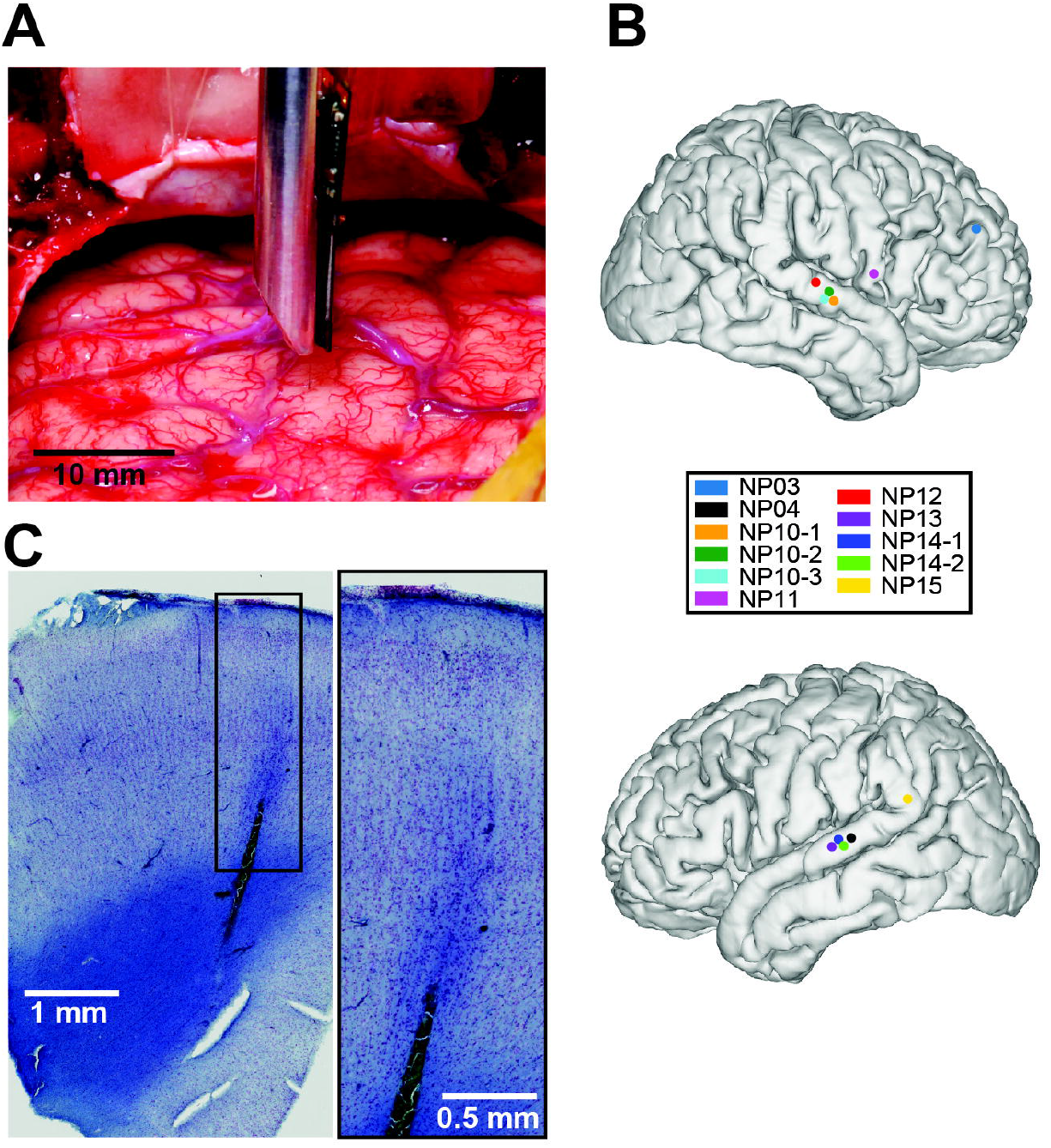
Intraoperative human recording with Neuropixels high-density silicon electrode arrays. A) Representative intraoperative photo after Neuropixel probe has been inserted in cortex (participant NP10). Inset scale bar corresponds to 10 mm. B) Insertion locations of Neuropixel probes with colors corresponding to participants, for all datasets with isolable single units. C) Representative histology of insertion tract lesion and neocortical cytoarchitecture, stained with luxol fast blue and neutral red. Inset with area of higher magnification. Scale bars below correspond to 1 mm (left) and 0.5 mm (right).

In cases where spiking was recorded, action potentials were apparent in the on-line filtered data (high-pass, 300Hz) within seconds of insertion (all 13 insertions with visible action potentials within 30 seconds of insertion upon post-hoc review of 300Hz high-pass filtered data across all 9 participants with spikes). In one case with visible spiking (NP06), the probe shank broke less than 1 minute after insertion and no putative single units were isolated from this brief recording. In one other insertion there was too much motion of the probe relative to the brain to isolate units (NP11, second insertion). In the 6 of 15 participants where no spiking was seen, two probes broke upon insertion, three had electrical noise which precluded successful recording (two participants awake, one anesthetized), and in one subject, no spiking was seen despite low noise and successful insertion (NP09). In all cases where the probe broke during or soon after insertion, participants were awake. As more datasets were collected, issues related to probe fracture or noise were mitigated (see Methods), with all 6 of the most recent recordings yielding spikes (Table 1). For the remainder of this paper, the 7 participants without isolable single units are excluded from further analyses.

While at least some identifiable spiking was present immediately after insertion, total spiking activity increased for several minutes after insertion. The data were clustered and curated resulting in putative single units that generally aligned well to the action potentials seen in the bandpass filtered data (Supplemental Fig. 2). Across all 11 insertions over 8 participants, half of recorded units became active within seconds after reaching target depth (8.8 seconds ± 9.2 seconds, mean ± standard deviation) and 75% of recorded units were active within 2 minutes (33 seconds ± 37 seconds). Recordings done in anesthetized patients took significantly longer for units to be active after reaching target depth (50% of units at least one spike, anesthetized 18.4 seconds ± 8.9 seconds, non-anesthetized 3.3 seconds ± 2.6 seconds, Wilcoxon rank-sum p=0.014; 75% of units at least one spike, anesthetized 67.3 seconds ± 39.8 seconds, non-anesthetized 13.5 seconds ± 12.8 seconds, Wilcoxon rank-sum p=0.038; Figure 2A). Note that participant NP03 was asleep but not anesthetized during recording and exhibits an empirical cumulative distribution function more similar to anesthetized patients (Figure 2A). Regardless, these timescales are compatible with meaningful recordings done within the time constraints of the intraoperative environment.

**Figure 2:**
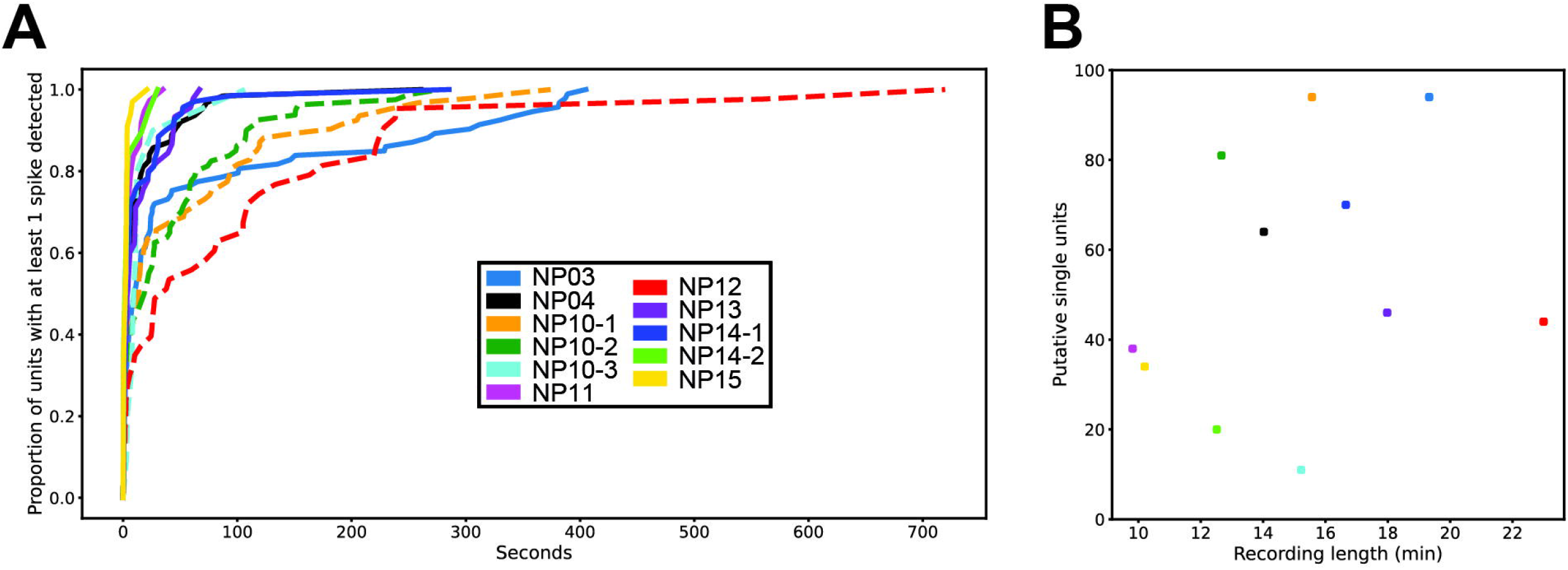
The majority of putative units are active within minutes. A) Empirical cumulative distribution functions of time of first spike for all putative single units shown for all insertions. X-axis, time in seconds. Time zero is after the probe is lowered to target recording depth. Y-axis, proportion of units that have at least 1 spike. Colors correspond to inset legend. Dashed lines represent recordings done in anesthetized patients. Note that NP03 was asleep but not anesthetized during recording. B) Recording length (x-axis, minutes) and number of putative single units recorded (y-axis). Colors correspond to legend inset in A).

There were 11 insertions that yielded putative single units. These recordings were 10 to 23 minutes in length (15 min 11 sec ± 3 min 47 sec, mean ± standard deviation). A total of 596 putative single units were identified, with 11 to 94 units per insertion and 34 to 94 simultaneously-recorded putative single units being isolated in each subject (best insertion per participant; 54.2 ± 27.2 putative single units, mean ± standard deviation across all recordings). There was no significant relationship between length of recording and putative single unit yield (Figure 2B, Spearman Correlation = 0.44, p = 0.17). There was no difference in putative single-unit yield between anesthetized and non-anesthetized states (Wilcoxon Rank-Sum p=0.78), and as expected, no difference was detected between hemisphere (Wilcoxon Rank-Sum p=0.47) or participant’s sex (Wilcoxon Rank-Sum p=0.65). There was no difference seen in single unit yield by recording location (superior temporal gyrus versus non-superior temporal gyrus; Wilcoxon Rank-Sum p = 0.92).

Clusters that passed manual curation spanned a range of quality across individual recordings (Supplemental Figure 3) as quantified by cluster metrics (see methods for more details; all metrics reported as median and interquartile range; spike amplitude: 136, 108 to 189μV; estimated proportion of amplitude histogram truncation: 0.0019, 0.0005 to 0.0266; signal-to-noise ratio: 4.2, 2.7 to 6.8; proportion of events violating 1ms inter-spike-interval: 0.0006, 0 to 0.0028; isolation: 0.993, 0.984 to 0.997; D-prime: 3.24, 2.77 to 3.86; firing rate: 0.91, 0.50 to 1.69 Hz; spike half-width, 0.33, 0.23 to 0.40ms; Figure 3).

**Figure 3:**
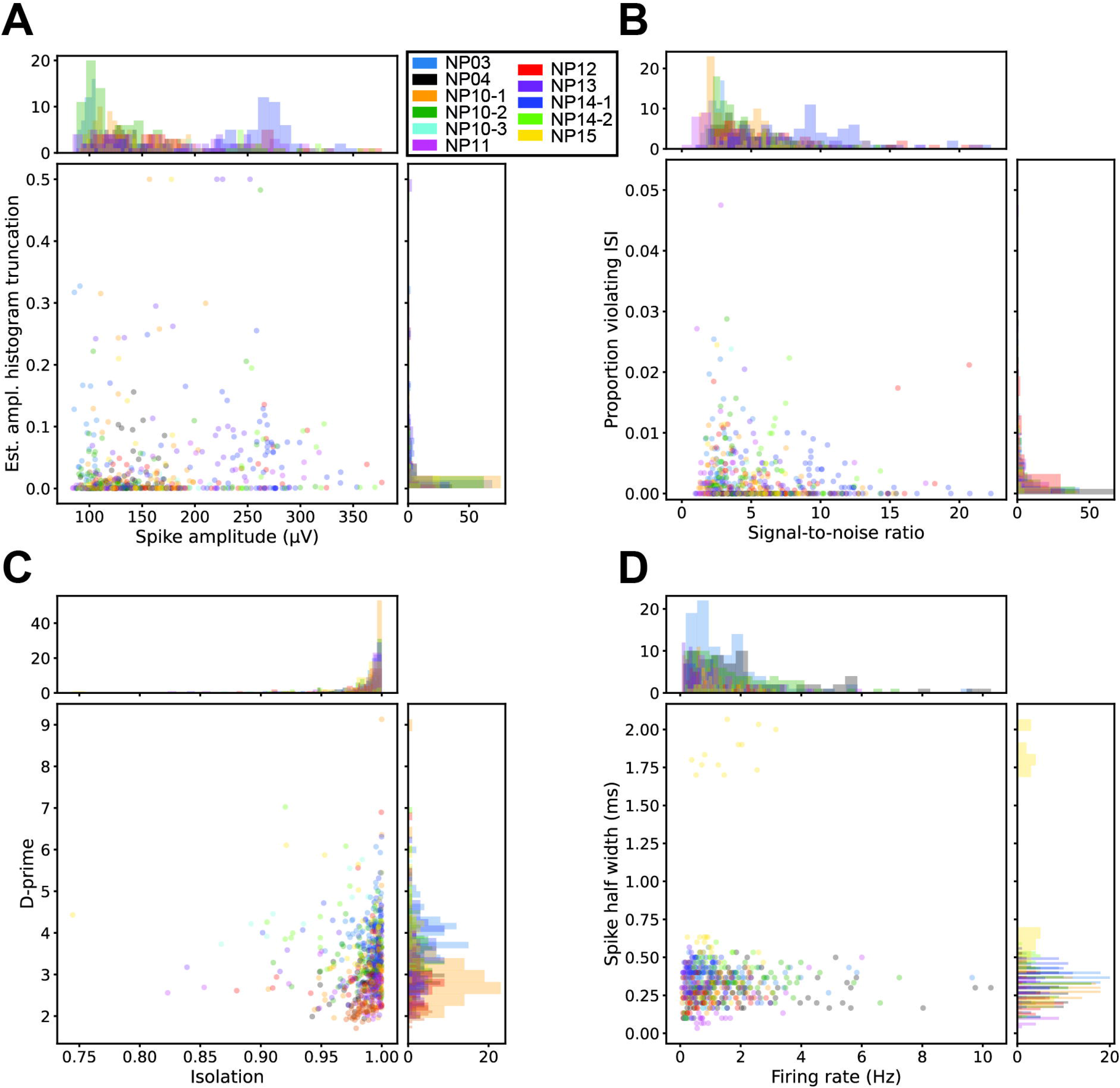
Neuropixels reliably isolate putative single units in human cortex. Cluster metrics plots for all 596 clusters from 11 datasets across 8 participants, legend inset. A) Scatter plot of amplitude (x-axis, μV) versus estimated proportion of amplitude histogram truncation (y-axis) B) as in A), but for signal-to-noise ratio (x-axis) and proportion of events violating the refractory period (inter-spike-interval less than 1 ms, y-axis). C) as in A), but for isolation (x-axis) and d-prime (y-axis). D) as in A), but for firing rate (x-axis, Hz), and spike half-width (y-axis, ms).

There are multiple types of relative motion between brain and recording electrodes, generally termed electrode drift. In the vertical axis of insertion (perpendicular to cortical surface), there is motion introduced from lowering the electrode array as well as from the physiologic pulsations from cardiac and respiratory cycles (Supplemental Video 1). Even in successful recordings, there was significant vertical motion that was only partially compensated by existing automated sorting methodologies and required manual curation. In the recording segments used for clustering, the size of pulsation at 10-20 cycles per minute, corresponding to the respiratory cycle, typically had vertical peak-to-peak amplitudes of 200μm. Pulsation occurring at 60-120 cycles per minute, corresponding to the cardiac cycle, typically had vertical peak-to-peak amplitudes of 100μm (Figure 4). In one insertion, the electrode drift associated with the pulsation resulted in data that did not yield putative single units (peak-to-peak amplitude 750μm, Supplemental Figure 4).

**Figure 4:**
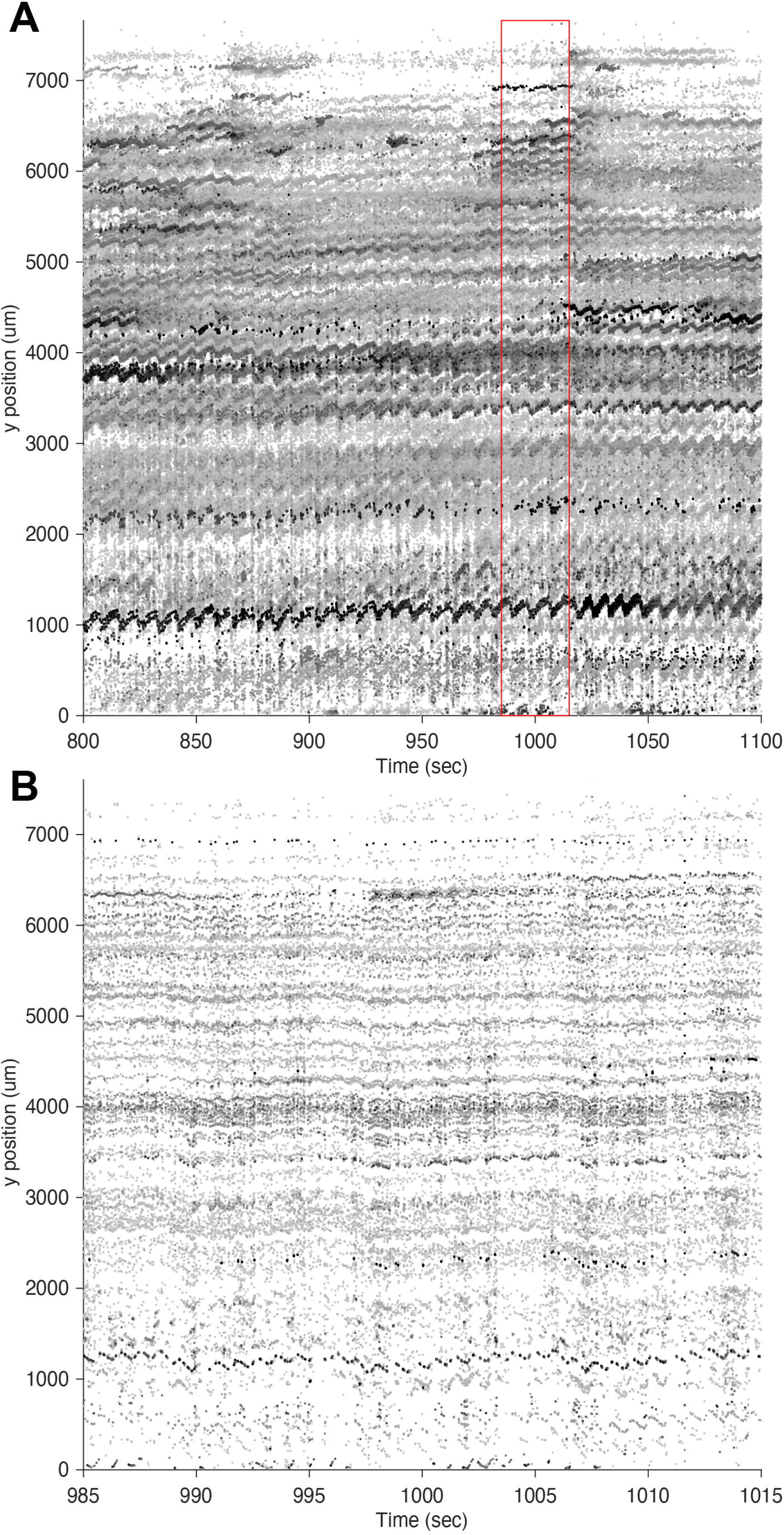
Electrode drift occurs predominantly in the vertical axis. Representative recording segment from participant NP03. A) Each dot represents a detected spike where relative darkness of greyscale reflects amplitude of spikes in arbitrary units (darker is higher amplitude, whitened and rescaled amplitudes, as in (Steinmetz et al., 2021). X-axis corresponds to time in seconds. Y-axis corresponds to spike position depth along the shank (0 μm is the deepest recording site). B) As in A) but for the sub-segment boxed in red in A), better demonstrating the timescale of cardiac cycle-associated pulsation.

Access to tens of simultaneously-recorded neurons across the depth of cortex allows analysis of the relative timings of the thousands of cell-pairs relative to their locations. Single-unit pairs’ relative activity was stratified by cell-pair coordination index (CCI, see methods). Briefly, cross-correlograms were calculated and the activity of cell pairs in close temporal proximity (within ± 50ms) was compared to the next 100ms period (−100 to -50ms and 50 to 100ms). Putative neuron pairs that fired more within ± 50ms than the next 100ms period had positive CCI; putative neuron pairs that fired more within the -100 to -50ms and 50 to 100ms time period, combined, than ± 50ms time period had negative CCI. Datasets with more than 1000 cell pairs were included in further analysis (Table 2). In 5 of the 8 datasets across 4 of 7 participants, cell-pairs with positive CCI were significantly closer together than those with negative CCI, 1 of the 8 datasets in one participant the converse was seen where cell pairs with positive CCI were significantly further apart, and in the remaining 2 of 8 datasets across 2 of 7 participants there was no relationship between distance between cell pairs and CCI (Figure 5, Supplemental Figure 5, Table 2). The simultaneous recording of tens of single-units and thousands of cell pairs across the depth of cortex allows for their relative spike timings and spatial locations to be studied at an unprecedented resolution in an awake human.

**Figure 5:**
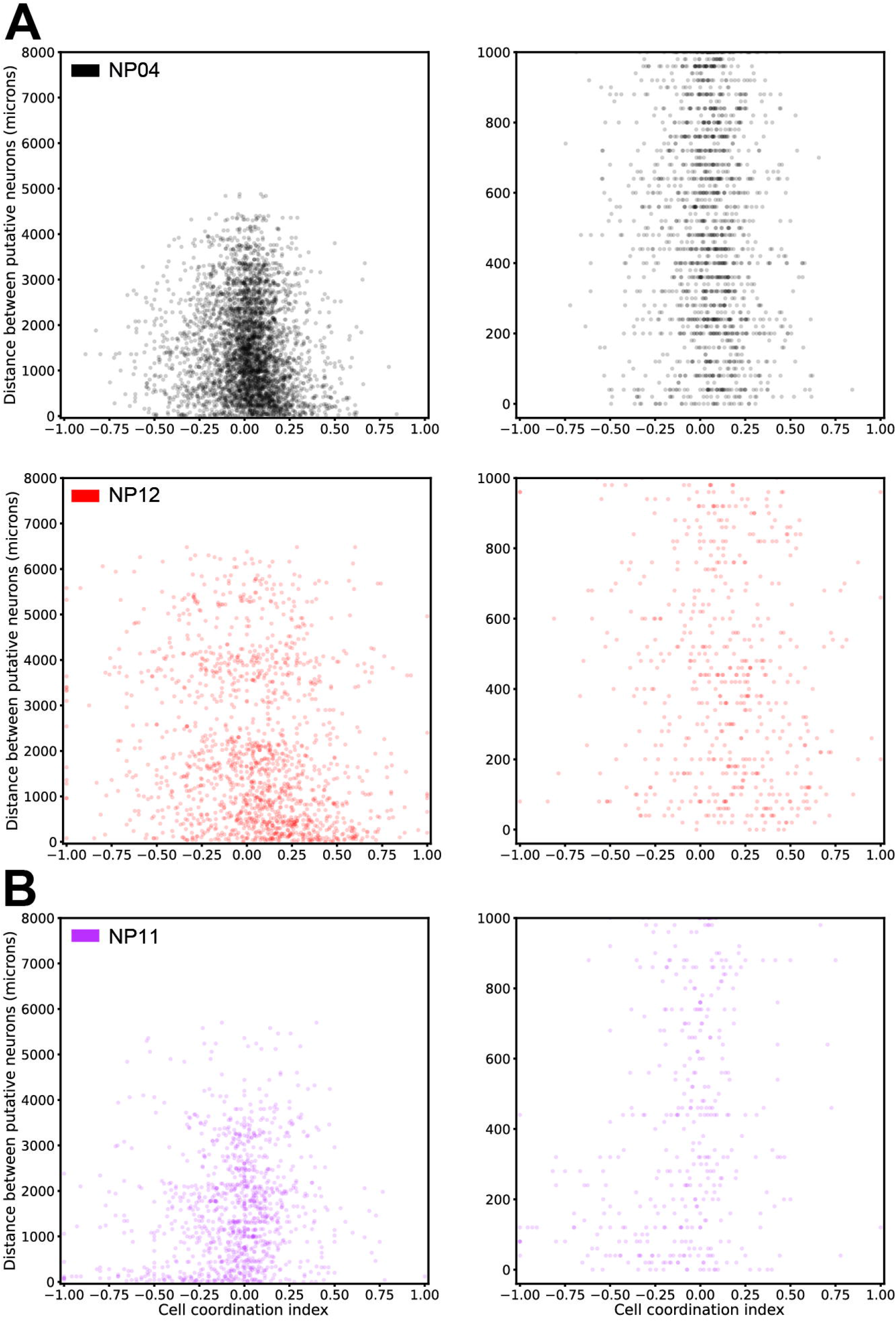
Cell-pair spike-timing differs by distance apart. A) Left, cell-pair coordination index (CCI, X-axis, see methods) by cell-pair distance apart (Y-axis, in μm), with identification inset top left, for 2 of the 5 datasets with positive CCI having significantly lower distances apart. Right, same as Left but only for cell-pair distances apart from 0 to 1000 μm. B) As in A), but for the dataset where negative CCI had significantly lower distances apart.

## Discussion

Large-scale recordings of single unit action potential firing across cortical layers provide unprecedented insight into the computational processes underlying neuronal function (Buzsáki, 2004). The ability to conduct these recordings in awake behaving human participants paves the way for studying the millisecond-timescale computations which underlie these behaviors. Towards that goal, we here provide a methodology capable of yielding dozens of simultaneously-recorded putative neurons within the constraints of the operative environment using the Neuropixels probe.

Here, we show 11 recordings that yielded a total of 596 putative single units in 8 of the initial 15 participants where Neuropixels recordings were attempted. Notably, failures related to intraoperative electrical noise and probe fracture were mitigated in later recordings. We note the key general goal of eliminating any alternating current device attached to the patient. Importantly, the operative table, electrocautery, and electronics associated with intravenous lines should be on direct current, including syringe pumps and blood/fluid warmers. Probe shank fractures were mitigated by either seeking locations with relatively thin pia or by performing piotomy.

Nearly all recordings that yielded putative single-units had visible spiking activity within seconds of insertion, and the majority of isolated units were active within a couple minutes of reaching target depth. In cases where the patient was anesthetized, there was a significantly longer latency before most putative single-units became active, though this delay was still on the timescale of minutes. Given the short recording length compared to animal experiments, this result may be heavily influenced by the firing rates of the isolated single units - there are likely insufficient spikes detected from low firing rate neurons to isolate them successfully using traditional spike-sorting algorithms. In addition to the inherent drawbacks imparted by short recording lengths, there is also the inherent increased variability in human participants, insertion sites, surgical exposures, and targeted regions which influence single-unit yield and quality.

All spike sorting here was done using Kilosort 2.5 with manual curation (Pachitariu et al., 2016) - necessary in large part due to the pulsation-associated electrode drift described below. While considerable variability is introduced from the operator biases inherent in manual curation in the spike sorting pipeline (Wood et al., 2004), the application of metrics (Buccino et al., 2020; Magland et al., 2020) allows objective assessment of quality. In this work, and for the first time in human high-density silicon array recording, cluster metrics for all putative single-units are reported, thereby establishing an objective point of comparison for quality and yield for human application of the Neuropixels probe.

Among the contributors to variability in recording quality, electrode drift from the pulsation of cardiac and respiratory cycles is a key component affecting quality. We found that in intubated patients, reducing positive end expiratory pressure helped reduce amplitude of both cardiac and respiratory cycles. There is a considerable amount of work necessary to counteract or compensate for brain movements both in physical and computational approaches. While Neuropixels 2.0 and associated motion correction algorithms (Steinmetz et al., 2021) show promise to better mitigate electrode drift in animal models, there is likely further computational work that will be necessary to address the larger-scale motions seen in human application.

In all cases in this work, a craniotomy was required for a range of clinical indications. Craniotomies expose a relatively large surface area of the brain to the air, thereby allowing access to a multitude of regions. While craniotomies can give access to nearly all surface brain structures and multiple structures simultaneously (and may allow for multiple simultaneous probe insertions in the future), this larger surgical exposure is associated with greater pulsation and brain movement relative to burr holes (which are typically 14mm to 25mm in diameter). The difference in exposure size is an important point of comparison to the simultaneously-completed work presented in (Paulk et al., 2021), where burr holes for deep brain stimulator implantation (giving access to dorsolateral prefrontal cortex) were used in 2 of the 3 successful recordings reported. In that work, the single case where they utilize a larger craniotomy exposure, they report 19 putative single units; here we show that the Neuropixels probe is capable of yielding up to 94 putative single units despite the technical challenges associated with larger surgical exposures.

Finally, we demonstrate the unprecedented capability to measure the millisecond dynamics across the depth of cortex in an awake, behaving human. As predicted from work in non-human primates across multiple regions (Constantinidis and Goldman-Rakic, 2002; Lee et al., 1998; Smith and Kohn, 2008), we report in 5 of the 8 datasets and in 4 of the 7 participants which met inclusion criterion (at least 1000 eligible cell pairs), in multiple regions, that cell pairs with a positive cell-pair coordination index (CCI, see methods) are significantly closer together than those with negative CCI. Though considerable work remains to be done to investigate these spatiotemporal relationships across different subjects, conditions, and locations, the application of Neuropixels probes in human intraoperative experiments provides unprecedented access to human neural processing.

### Single-unit recordings in humans

Single unit recordings in human brain have paved the way for exceptional findings, including ‘concept cells’ (Quiroga et al., 2005), tremor-related cells (Levy et al., 2000), place cells (Ekstrom et al., 2003), mechanisms underlying seizure generation (Wyler et al., 1982), and anatomical subdivisions of language and memory processing (Ojemann et al., 1988, 2002). The earliest single unit recordings in human brain were conducted in the 1950s (Ward and Thomas, 1955). For decades, these recordings were conducted with a single microwire, tungsten electrode (Bertrand and Jasper, 1965), or a small bundle of microwires, typically in patients with seizures or for confirmation of anatomical targeting in movement disorders (Benazzouz et al., 2002; Hutchison et al., 1998). While valuable data to inform neuronal processing, the unit yields were very low (e.g. (74 units across 9 patients) (Fried et al., 1997); 90 units across 17 patients (Wyler et al., 1982)) and localization of individual microwire placement was difficult or impossible.

More recent advances in human single unit recordings include use of the Utah array. Spiking activity from the Utah array has successfully been used by a tetraplegic participant to control a cursor-based brain machine interface (BMI) (Hochberg et al., 2006) and a 17-degree-of-freedom robotic limb (Aflalo et al., 2015). Gradually degrading but viable signal has been recorded from Utah arrays for up to 1500 days (Hughes et al., 2021); however, substantial immune response and tissue reorganization at the implant site is also common (Szymanski et al., 2021). The Utah array samples units across an X-Y plane and is not able to record units across the depth. Another class of probes includes microelectrode arrays, such as the Michigan array. These arrays are silicon-based probes manufactured using electron-beam lithographic techniques (Choi et al., 2018; Pochay et al., 1979); these arrays allow for recording across the depth, but their use has been mainly limited to animal models.

For the microwire and Utah array approaches discussed above, a given unit can only be detected on one recording channel, as the shanks/contacts are spaced at least 100um apart (Harris et al., 2016). Some microelectrode arrays have contacts less than 50um apart, allowing for a given unit to be detected on multiple channels. This increases not only the number of recording contacts, but also the accuracy of spike sorting because the spatiotemporal profile of a given unit can be better differentiated from other units when acquired from multiple recording channels (Rossant et al., 2016). The Neuropixels probe extends this even further, with a larger number of densely-spaced contacts.

### Neuropixels probe

The major advance demonstrated here is recording in human brain with densely-spaced high channel count simultaneously acquired across the cortical thickness. This is directly enabled by the Neuropixels probe. The Neuropixels probe is one of a handful of advanced neural recording probes which have built on CMOS-based microelectrode silicon probes (Hong and Lieber, 2019). The Neuropixels probe contains 960 contacts arranged in a 4 column checkerboard pattern; select subsets of 384 contacts can be recorded from simultaneously (Jun et al., 2017). We recorded in a ‘long column’ configuration, in which one contact was used at each depth, and adjacent contacts alternated between two columns. This geometry of contacts allowed for semi-automatic spike sorting.

Kilosort is an automated spike-sorting algorithm that has been designed to directly leverage the geometry of Neuropixels probes (Pachitariu et al., 2016; Steinmetz et al., 2021). We used version 2.5 of the software which includes more advanced drift correction. Because contacts are arranged with 20um spacing, waveforms for one unit are typically detected on multiple adjacent contacts. Kilosort employs a clustering algorithm across channels in order to separate units from different putative neurons using template waveforms. While unsupervised performance of the algorithm exceeded that of prior spike sorting methods, manual review and curation was conducted using the Phy graphical interface (Rossant et al., 2016).

### Procedural hurdles for Neuropixels in humans

The translation of Neuropixels technology from animal models to human participants required several procedural and experimental hurdles to be overcome. First and foremost is the health and safety of human participants. Therefore, risk related to insertion of the Neuropixels probe and any modifications to standard procedure in the operating room must remain at a minimum. While the thicker shank of Neuropixels 1.0 NHP-short probes were more robust against movement in the human brain, shank breakage did still occur in some experiments. In each instance, the broken shank was retrieved. Even in cases where the shank remains intact, microscopic tissue damage occurs along the insertion trajectory. For both these reasons, we only inserted the probe in tissue that was planned for resection in the same procedure.

To minimize the risk of infection, all probes and accessories were sterilized using standard medical-grade processes. The sterile surgical field was maintained and extended to part way along the length of the cable connecting the headstage to the Neuropixels PXIe Acquisition Module. A gowned and gloved surgeon connected the headstage to the cable and probe and positioned the probe in the micromanipulator. No probes were re-used across participants.

### Experimental hurdles for Neuropixels in humans

The intraoperative environment imposes limitations on experimental recordings. When Neuropixels are used in animal models, the probe is typically inserted quite slowly (e.g. 2-10um/second) (Böhm and Lee, 2020; Kostadinov et al., 2019; Park et al., 2019; Schröder et al., 2020) in order to optimize recording quality (Fiáth et al., 2019), and then allowed to sit for 5-60 minutes prior to starting experimental data acquisition. However, to minimize added time in the operating room and under anesthesia, we restricted recordings to ∼20 minutes. Therefore, we started acquiring data minutes after probe insertion. Future work (likely in animal models) will be required to understand how settling time and potential stunning effects from insertion alter spike rates or patterns. Our total recording duration was also shorter than may be optimal for spike sorting, leading to higher noise clusters. Nevertheless, we were able to acquire tens of units within minutes of probe insertion and identify distinct clusters.

The operating room is also an electrically noisy environment. During recordings, we turned off and unplugged all non-essential powered equipment. Equipment that contributed the most noise included anesthesia equipment, neuromonitoring, and the IR sensor from neuronavigation. We found that using separate needles for ground and reference reduced noise rather than shorting these together. Despite these de-noising steps, the noise levels remained higher than can be achieved using a Faraday cage for animal experiments.

A major challenge for recording single units in human cortex is the movement of the brain caused by pulsation. The skull is secured in a clamp, and the micromanipulator apparatus holding the Neuropixels probe was secured relative to this clamp. However, the brain movement relative to the skull was reflected in recordings. These ‘movement artifacts’ can be reflected as a given putative unit shifting up and down on the contacts on the shank, or slight variation in the detected waveform. Future iterations of recording procedure may benefit from untethering the probe from the insertion micromanipulator apparatus, to ideally allow the inserted probe to move with the brain. Our initial attempts to manually correct for movement based on video recordings or frequency of pulsation were not successful – while heart rate was recorded, the brain movement was not stereotyped based on heart rate, and there were multiple axes of movement (primarily vertical relative to the probe, but also small amounts of lateral movement). Additional computational methods to model and compensate for the pulsation-associated electrode drift is ongoing but faces two key challenges in human intraoperative experiments. First, the amplitude of movement is far larger than in animals, and at least an order of magnitude larger relative to rodents. Second, short recording times reduce the number of recorded spikes for every neuron. Together, this means that fewer spikes are spread across a larger region of the array, making it far more difficult to correct for the motion. Densely packed and linearly arranged (in the axis of the motion) columns of electrodes, such as in the Neuropixels 2.0 design (Steinmetz et al., 2021), may be much more capable to compensate for the motion seen in human participants. Despite these challenges, we conducted post-hoc algorithmic motion correction using Kilosort 2.5 as described above.

As with any study, there are shortcomings and weaknesses in this method. While the intraoperative environment provides access to recording single units in human cortex, the duration of recordings and possible tasks and behaviors are limited (e.g. only non-ambulatory). Sub-chronic implant of a Neuropixels probe or other comparable technology would allow for a richer dataset to explore the longitudinal behavior of unit firing. However, this is accompanied by considerable barriers regarding material safety, stabilization, and signal integrity. The current geometry of Neuropixels probes also precludes recordings in deeper subcortical structures. Even if probes were made longer, there remains the risk for probe breakage and the need to retrieve the shank. Currently available Neuropixels probes do not include the capacity for electrical stimulation. Future iterations may add this feature, at which point a new set of experiments looking at activation of the microcircuit across cortical layers will be possible. And as with any study incorporating spike sorting, there are false positive and false negative errors in assigning spiking to individual units/clusters (Harris et al., 2016).

In summary, use of the Neuropixels probe in humans offers widespread access to large ensembles of cortical neurons across the depth of the neocortex. While there are practical and technical limitations to experiments that can be conducted in the intraoperative environment, there are still myriad insights into human behavior that can be gained using this approach.

## Figure and Table Legends

**<u>Table 1: Participant information</u>**

Participants are listed in chronological order. The majority had surgery for medically refractory epilepsy. Note that participant NP03 was asleep but not anesthetized during recording.

**<u>Table 2: Cell-pair coordination index and distance</u>**

Participants and insertions listed in chronological order. Datasets with more than 1000 cell pairs were included in further analysis. All distances in microns. P-values for Wilcoxon rank-sum.

## Supplemental Figure and Movie Legends

**Supplemental Figure 1:**
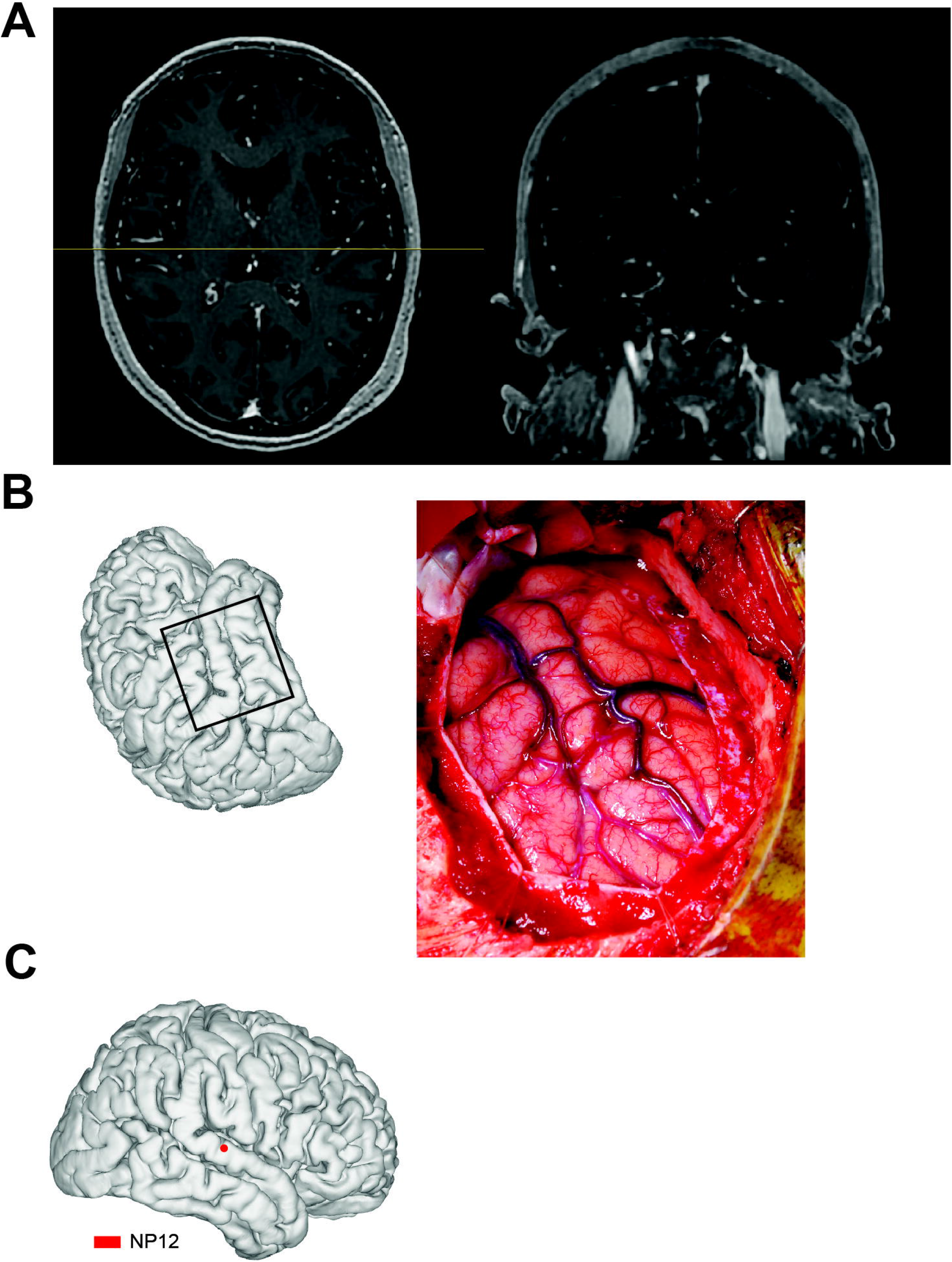
Localization of insertion location. Example probe localization for NP12 where Neuropixels recording was targeted to the right superior temporal gyrus. A) Preoperative MRI showing axial reformat (left, slice thickness = 1.0 mm), and coronal reformat (right, specific location marked by yellow line on axial reformat, slice thickness = 1.0 mm). B) Left, schematic depicting orientation of patient and approximate craniotomy exposure (boxed). Right, intraoperative photo. C) 3-dimensional reconstruction of brain and electrode locations using imaging in A) and image in B), with the red dot showing estimated insertion location.

**Supplemental Figure 2:**
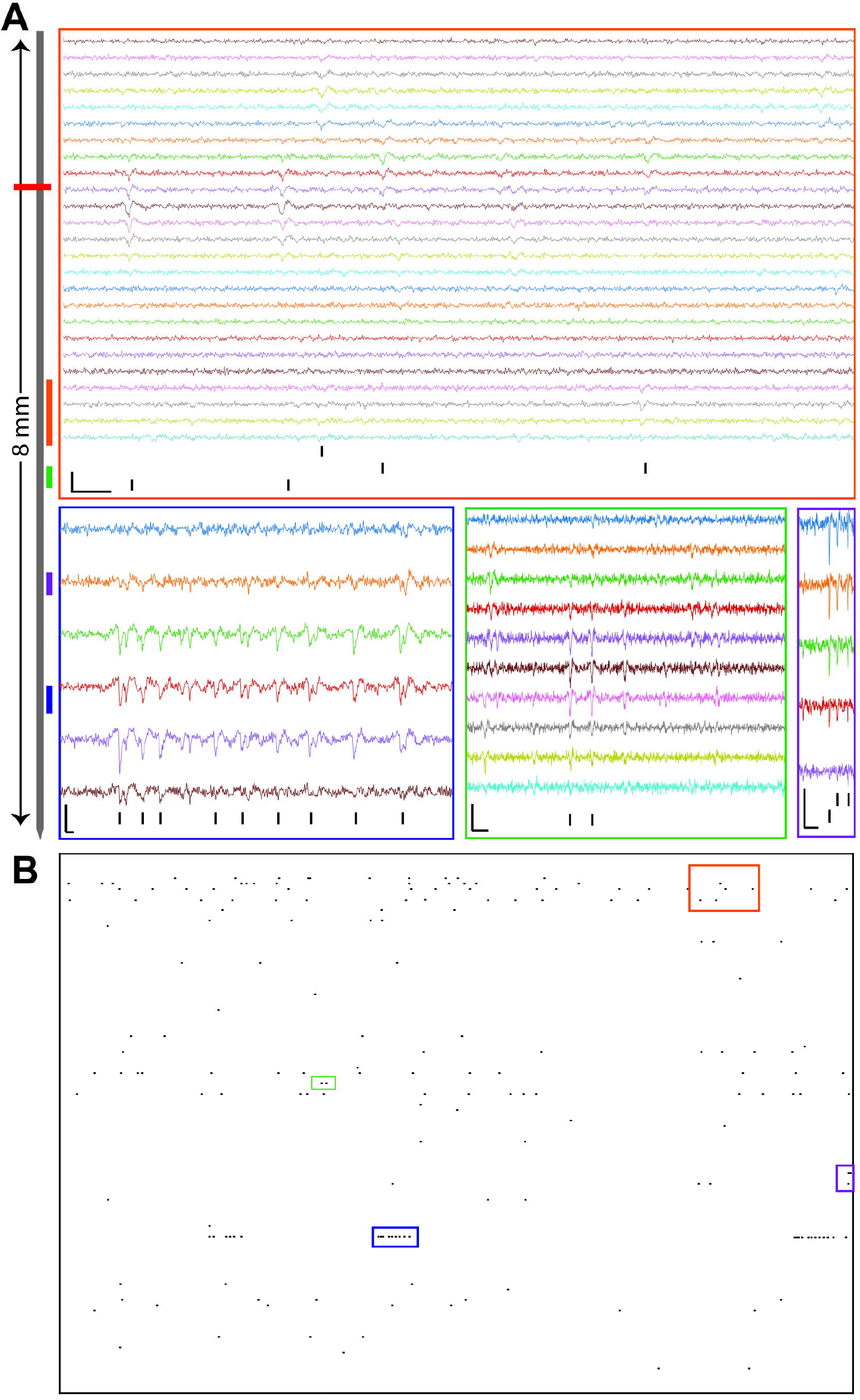
Neuropixels produce high-quality intraoperative recording in human cortex. Representative recording snippets from NP03, filtered from 300 to 6000 Hz. A) Left, schematic of electrode array. Horizontal red line represents estimated brain surface depth. Total length of the probe shown corresponds to 8 mm, with 6.8 mm below the brain surface. Recording excerpts, boxed, from channels at the color-corresponding depths, with rasters for the 1-3 units represented in the excerpt. Inset scale bar in bottom left corresponds to 200 μv and 5 ms. B) Spike rasters for 94 putative units following spike sorting and manual curation in 1 second of data. Each row is a raster from a putative single unit. Note not all 94 units are active in this second of data.

**Supplemental Figure 3:**
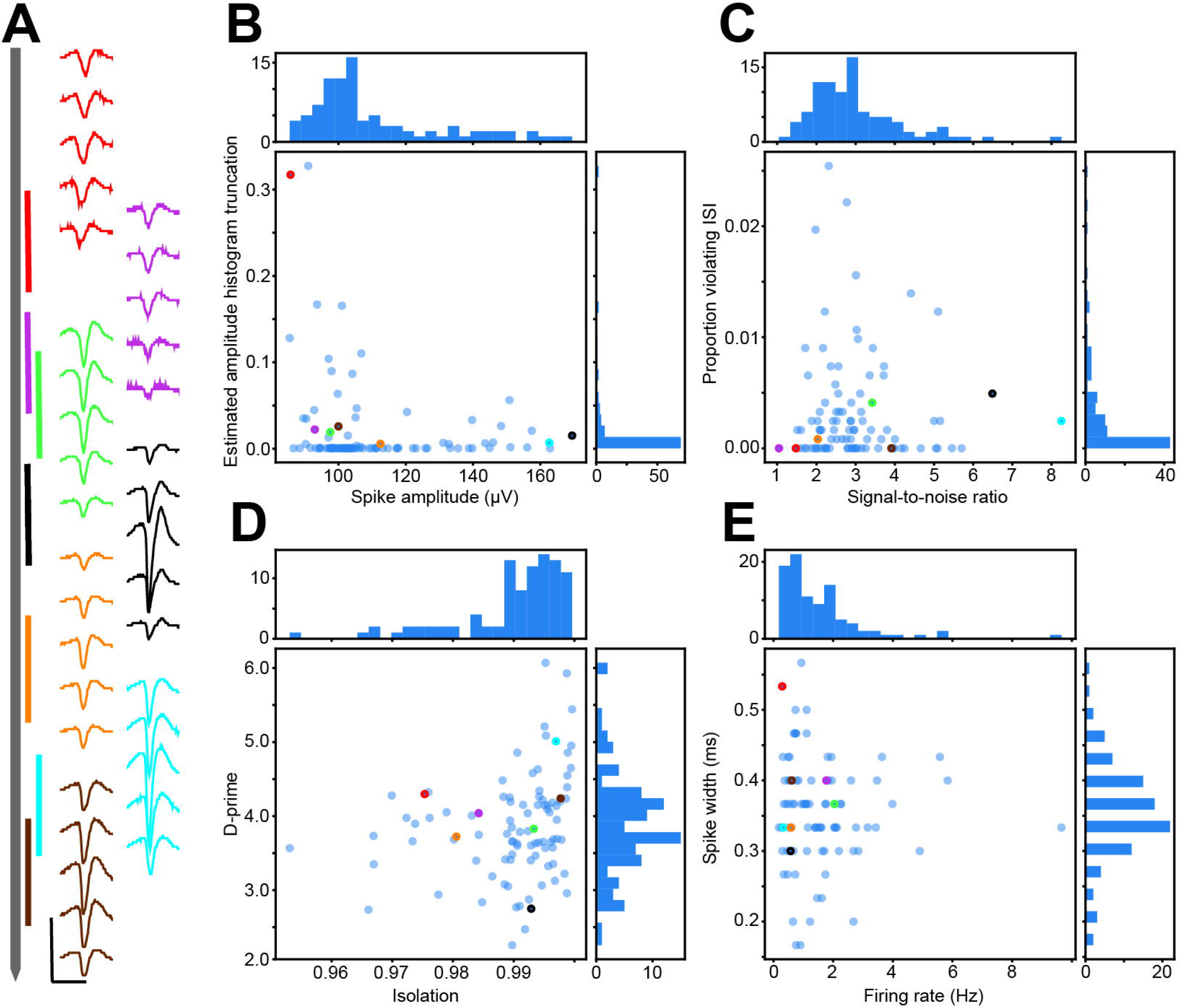
Diversity of single unit quality. A) Clusters from participant NP03. Left, schematic of probe shank with color-coded vertical lines indicating depth of example clusters. Example clusters selected to represent diversity of cluster metrics. Total length of the shank shown is 8 mm. Right, average waveforms of example clusters. Scale bar at bottom corresponds to 150 μV and 2.5 ms. B) Scatter plot of amplitude (x-axis, μV) versus estimated proportion of amplitude histogram truncation (y-axis) for all 94 clusters from participant NP03, with color-corresponding example clusters and axes-corresponding histograms. C) as in B), but for signal-to-noise ratio (x-axis) and proportion of events violating the refractory period (inter-spike-interval less than 1 ms, y-axis). D) as in B), but for isolation (x-axis) and d-prime (y-axis). E) as in B), but for firing rate (x-axis, Hz), and spike half-width (y-axis, ms).

**Supplemental Figure 4:**
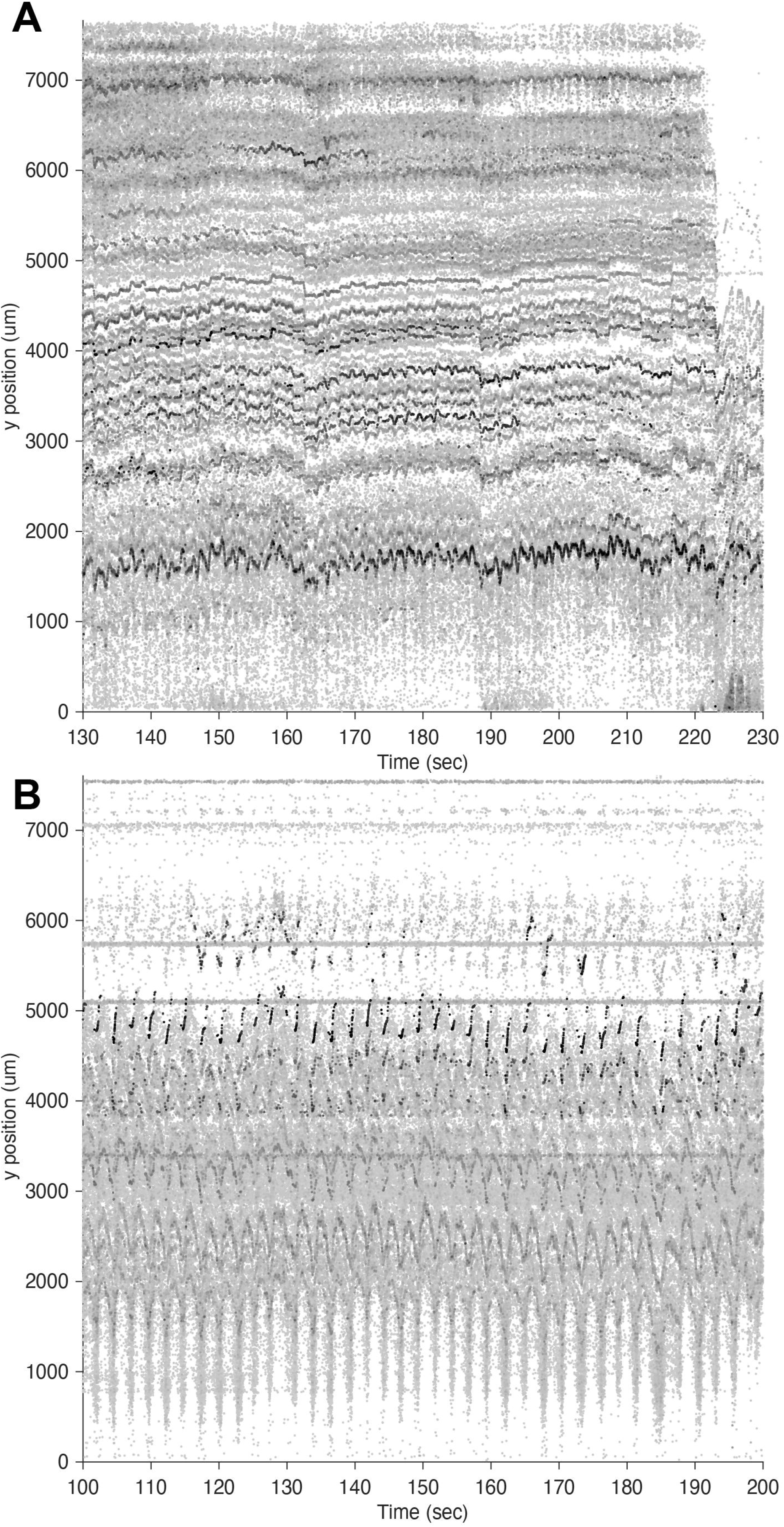
Physiologic pulsation can cause overwhelming drift. Representative recording from two different insertions in participant NP11. A) Recording segment which was used for spike sorting and yielded putative single units. Each dot represents a detected spike where relative darkness of greyscale reflects amplitude of spikes in arbitrary units (darker is higher amplitude, whitened and rescaled amplitudes, as in Steinmetz et al. 2021). X-axis corresponds to time in seconds. Y-axis corresponds to spike position depth along the shank (0 μm is at tip). Shank is withdrawn from the brain around 223 seconds. B) Recording segment that did not yield any putative single units due to overwhelming electrode drift.

**Supplemental Figure 5:**
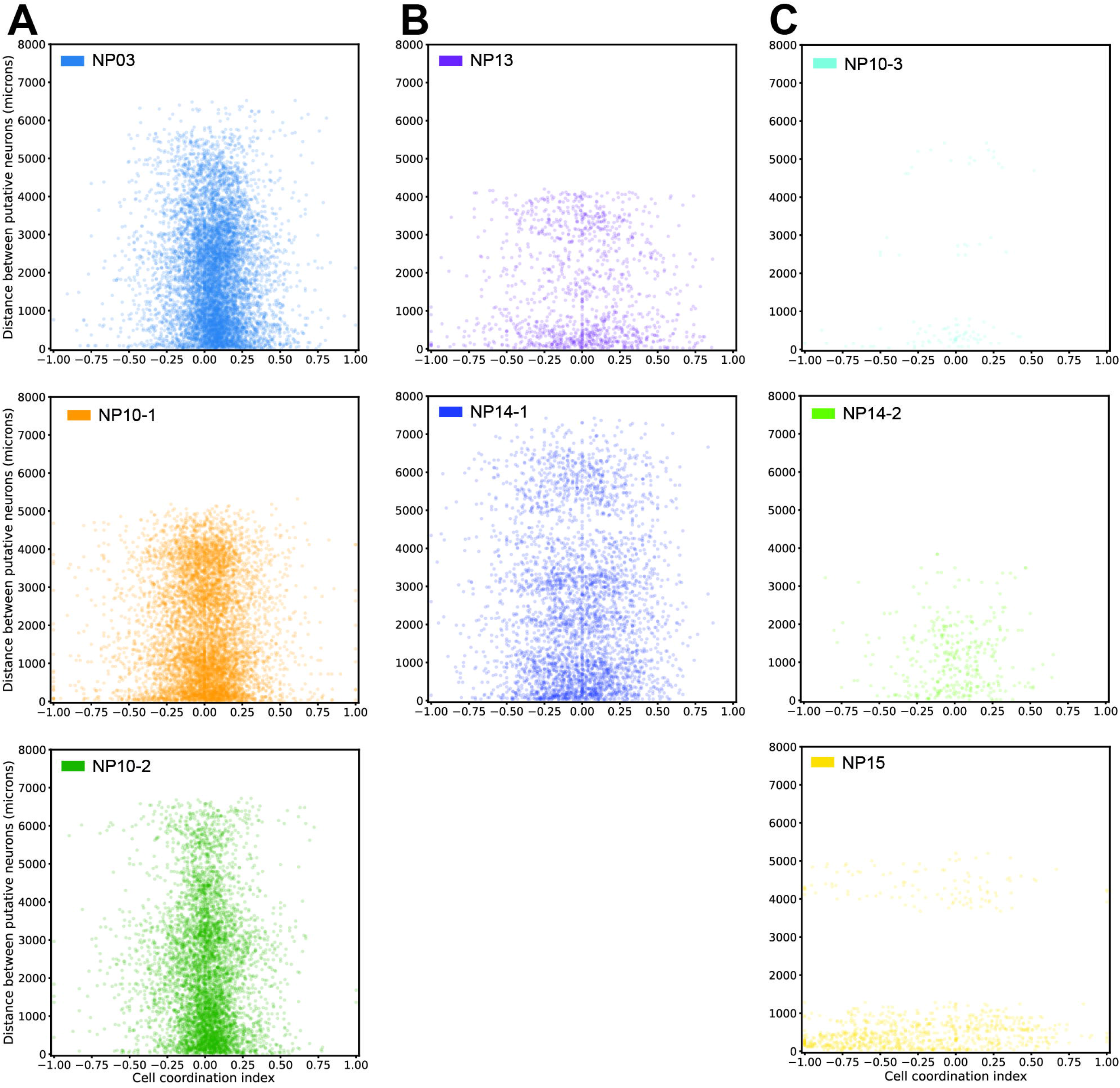
Cell-pair spike-timing by distance apart. A) Cell-pair coordination index (CCI, X-axis, see methods) by cell-pair distance apart (Y-axis, in μm), with identification inset top left, for the 3 of 5 datasets with positive CCI having significantly lower distances apart. B) As in A), but for the 2 datasets where there was no relationship between CCI cell-pair distance. C) As in A) and B), but for the 3 datasets that were excluded for having less than 1000 cell pairs.

**<u>Supplemental Movie 1: Insertion and intraoperative recording</u>**

Intraoperative video of an insertion for participant NP12 demonstrating a representative deformation associated with penetration of pia, and pulsation associated with the cardiac and respiratory cycles.

## Notes

### Competing Interest Statement

The authors have declared no competing interest.

